# Full-mouth photoacoustic/ultrasound imaging of the periodontal pocket with a compact intraoral transducer

**DOI:** 10.1101/2022.03.31.486608

**Authors:** Lei Fu, Reza Khazaeinezhad, Ali Hariri, Baiyan Qi, Casey Chen, Jesse V. Jokerst

## Abstract

Periodontitis is a public issue and imaging periodontal pocket is important to evaluate periodontitis. Regular linear transducers have limitations in imaging the posterior teeth due to their geometry restrictions. Here we characterized a transducer that can image the entire human mouth including assessment of periodontal pockets via a combination of photoacoustic and ultrasound imaging. Unlike conventional transducer design, this device has a toothbrush-shaped form factor with a side-view transducer to image molars (total size: 1 × 1.9 cm). A laser diode was integrated as the light source to reduce the cost and size and facilitates clinical transition. The *in vivo* imaging of a molar of a periodontal patient demonstrated that the transducer could image in the posterior area of gum *in vivo*; the value determined by imaging was within 7% of the value measured clinically.

## 1. Introduction

Periodontitis is a common disease caused by subgingival bacteria that destroy the supporting structures of the teeth^1^. Deeper periodontal probing depths and gingival inflammation are common features of periodontitis^2^. Oral health professionals measure probing depths with a metallic probe placed in the pocket between the gingiva and tooth^3^. However, periodontal probing is subjective to the probing force, probe angulation, and the insertion point^4, 5^. Periodontal probing can also cause bleeding and is painful to the patient and time-consuming for the provider.

We previously reported photoacoustic/ultrasound imaging to image the periodontal pocket in an *ex vivo* swine model and *in vivo* human subjects^6, 7^. However, only anterior human teeth could be imaged because of the bulky geometry of the transducer. The handle of regular linear transducer needs to project out orthogonally from the tissue surface that is being imaged, which makes the transducer cumbersome to operate in the posterior area of gum. Unfortunately, most periodontitis occurs on posterior teeth including molars and pre-molars. Although there are clinical and small transducers e.g., endoscopic transducers that could be potentially adapted for periodontal imaging^8^, they usually have low central frequency at around 5 MHz, which is insufficient to resolve small structures of tooth^9, 10^. For this reason, it is important to develop a photoacoustic/ultrasound transducer that can image posterior teeth with high resolution. Concurrently to this, we were motivated to use a light source that is rugged and affordable. While our prior work used a Q-switched laser, such lasers are very expensive and bulky^6, 7^. In recently years, many groups have developed laser-diode or LED-based photoacoustic imaging system which is more portable and affordable than Q-switch laser^11^.

In this work, we characterized a compact and affordable photoacoustic/ultrasound transducer for *in vivo* full-mouth periodontal pocket imaging. The transducer has a form factor reminiscent of a toothbrush and thus can image all the human teeth *in vivo*. It integrates a 19-MHz ultrasound transducer with optical modules for photoacoustic imaging. We reduced the cost of the system by using laser diode as light source. Here, we first describe the transducer and considerations of using it to image posterior teeth. We then characterized the performance of the whole system in photoacoustic imaging. Finally, we imaged swine teeth *ex vivo* and a molar of a periodontal patient *in vivo*.

## 2. Materials and Methods

### 2.1 Hardware

We used a high-frequency ultrasound transducer (SS-19-128, StyloSonic, USA) in a custom handpiece. It has central frequency of 19 MHz and average -6dB bandwidth of 48.9%. The transducer has 128 elements with element pitch of 78 μm. The laser diode generates laser pulses in 1 kHz in 808 nm. The pulse energy is 0.7 mJ/cm^2^ and the pulse width is 100 ns. A research ultrasound data acquisition system (Vantage; Verasonics, Inc., Kirkland, WA, USA) was used to receive, process, and reconstruct the photoacoustic/ultrasound signals. The Vantage system has 256 channels with maximum sampling rate of 62.5 MHz. It provides output triggers to synchronize with the laser diode.

### 2.2. Characterization

Characterizations of photoacoustic imaging used a tissue-mimicking phantom made by 20% intralipid solution. We used pencil leads (0.2 mm diameter) to evaluate light homogeneity. We used a photodetector (PE50-DIF-C, Ophir Photonics Corporation) connected with an oscilloscope (SDS 1202X-E, Siglent Inc.) to measure the light stability. Three polyethylene tubes filled with cuttlefish ink were used to evaluate the imaging depth of the transducer. A human hair (0.1 mm diameter) was used to evaluate the resolution of photoacoustic imaging. A nichrome wire (0.03 mm diameter) was used to evaluate the resolution in ultrasound imaging.

### 2.2 Swine teeth

Swine jaws were obtained from a local abattoir. One swine tooth (1^st^ molar) was prepared for the ultrasound imaging in **Fig. 3**, and two more swine teeth (1^st^ molar and 1^st^ pre-molar) were prepared for photoacoustic/ultrasound imaging in **Fig. 4**.

### 2.3 Contrast agent

The melanin nanoparticles in cuttlefish ink (Nortindal, Spain) were used as a photoacoustic contrast agent to highlight the pocket as described previously^6, 7^.

### 2.4 Periodontal pocket imaging

The photoacoustic images were collected along with ultrasound data for anatomy. The photoacoustic image shows the distribution of the contrast agent in the pocket and was used to measure pocket depth. Pocket depth measurements were measured by Marquis probing method and photoacoustic method. The Marquis probe has black/white rings marking the probing depth. The probe was placed into the pocket along the direction of tooth root^12^ for conventional probing assessment.

### 2.5. Participant

All work with human subjects was approved by the University of California – San Diego (UCSD) and University of Southern California (USC) IRBs and conducted according to the ethical standards set forth by the IRB and the Helsinki Declaration of 1975. The participant gave written informed consent and teeth were imaged non-invasively. All subjects were >18 years old and able to provide consent. One healthy subject and one periodontal patient were recruited for this study. The imaging of periodontal patient was handled by a board-certified periodontist (author CC) at USC dental school.

## 3. Results and Discussion

We first describe the device (**Fig. 1**) and characterized the performance of the transducer in photoacoustic imaging (**Fig. 2)**, and then validate its performance in ultrasound imaging (**Fig. 3**). We eventually performed both *ex vivo* and *in vivo* periodontal pocket imaging (**Fig. 4**).

**Fig. 1.**
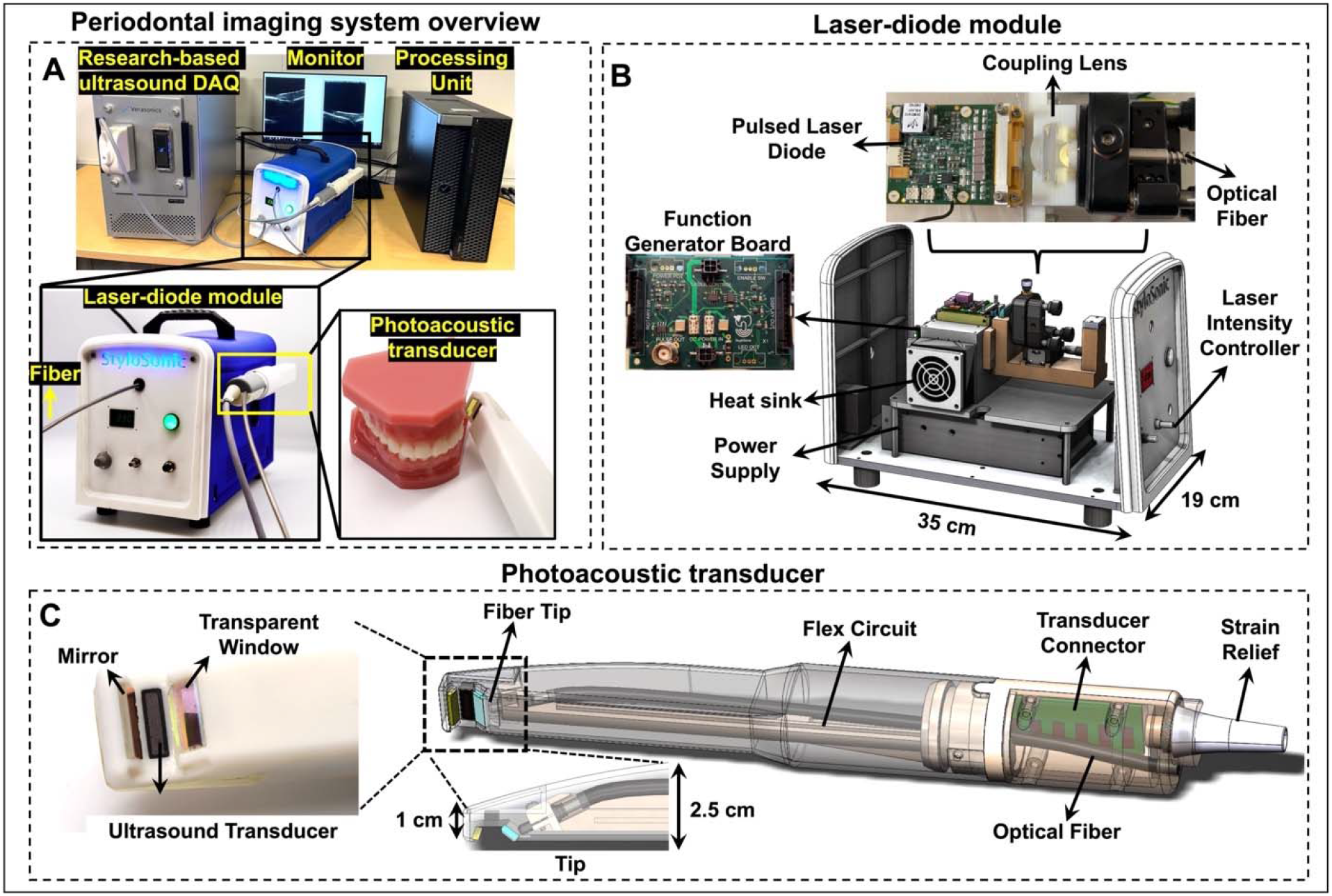
Full-mouth periodontal imaging system. A) System overview. A laser-diode module is the light source and light pulses are delivered to the transducer via a fiber. The transducer is integrated into a handheld device. A research-based ultrasound DAQ system processed and reconstructed the photoacoustic/ultrasound signals. B) A laser-diode module houses the pulsed laser diode, coupling lens, customized function generator board, heat sink, and power supply. The front panel has the control button to adjust the laser intensity. C) The photoacoustic transducer includes an ultrasound transducer, transducer connector, flex circuit, an optical fiber, transparent window, and mirror. The small tip of the handpiece (10 mm×19 mm) facilities full-mouth scanning.

**Fig. 2.**
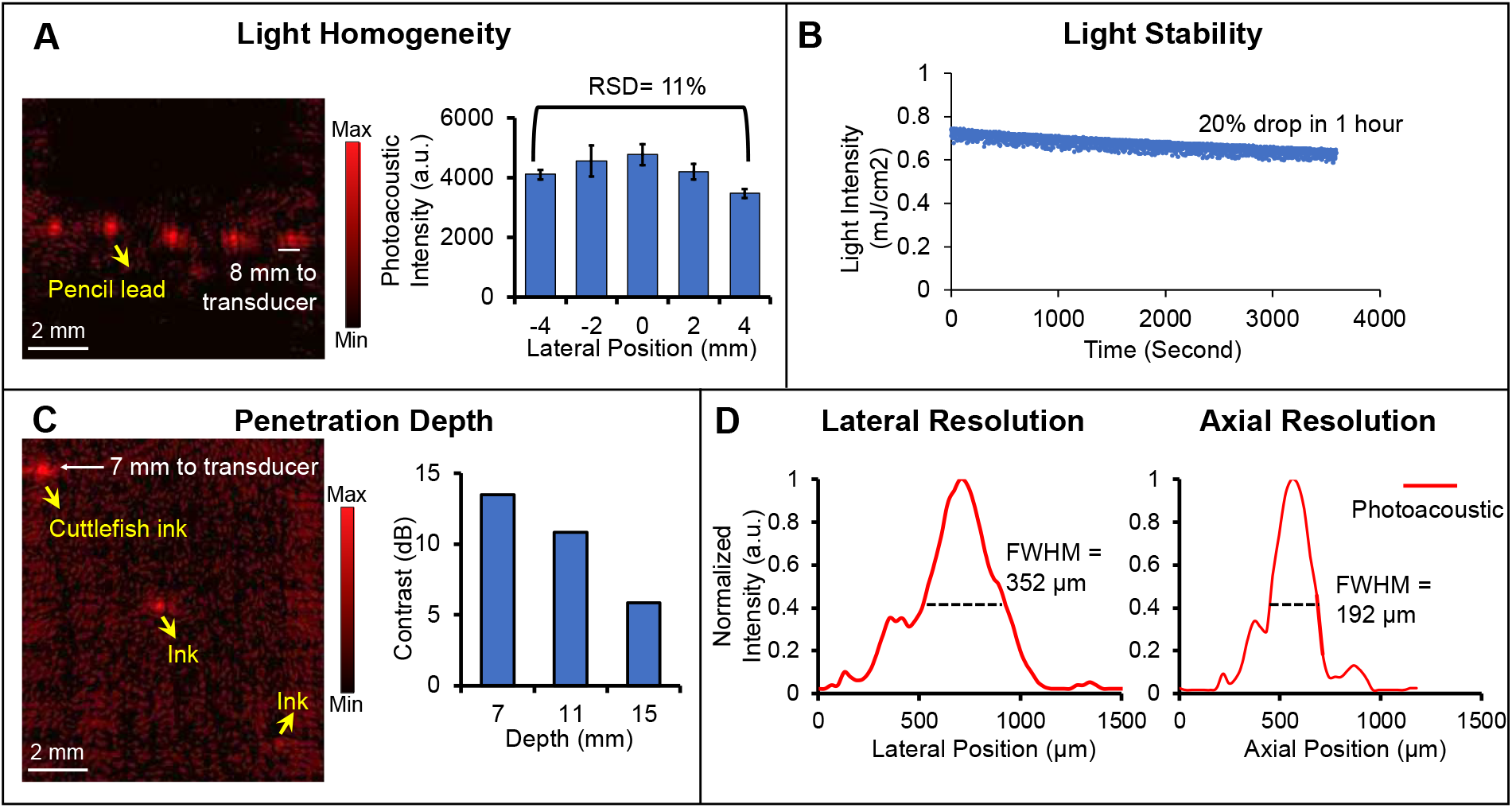
Performance characterization of the periodontal imaging transducer. A) Light homogeneity evaluation. Photoacoustic imaging of five pencil leads placed parallel over 10 mm and 8 mm under the transducer. The chart shows the relative standard deviation (RSD) of the photoacoustic intensity of the five pencil leads. B) Laser-diode power stability versus time in 1-hour scale. C) Imaging depth evaluation. Three tubes containing the contrast agent are put in a tissue mimic phantom located at different depths of 7 mm, 11 mm, and 15 mm from the transducer. The right chart shows the SNRs of the three tubes. D) The lateral and axial photoacoustic amplitude distributions along a 100-*μ*m hair: 352 µm and 192 µm are the lateral and axial resolution in photoacoustic mode, respectively.

**Fig. 3.**
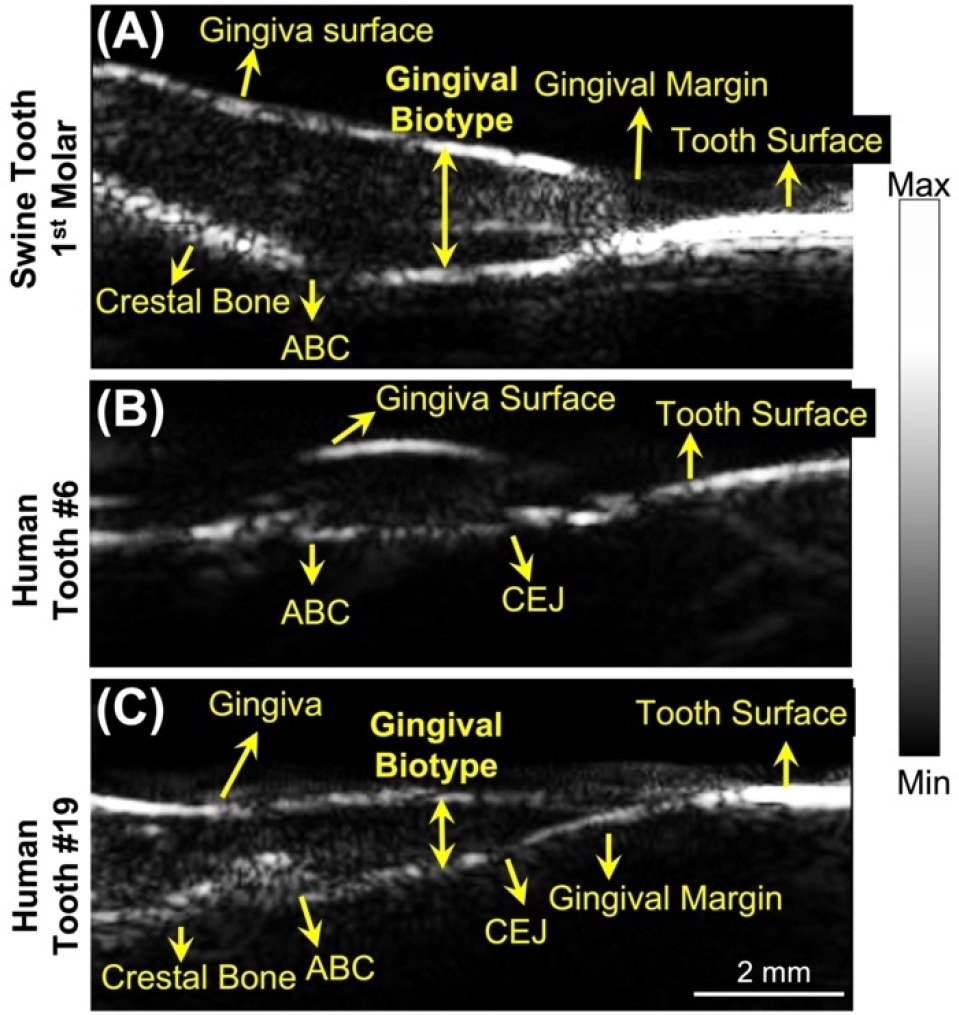
Ultrasound imaging of swine and human teeth. A-C) are ultrasound images of a swine 1^st^ pre-molar (A), a human cuspid #6 (B), and a human 1^st^ molar #19 (C). All the images are in sagittal view. Alveolar bone crest (ABC). Cementoenamel junction (CEJ).

**Fig. 4.**
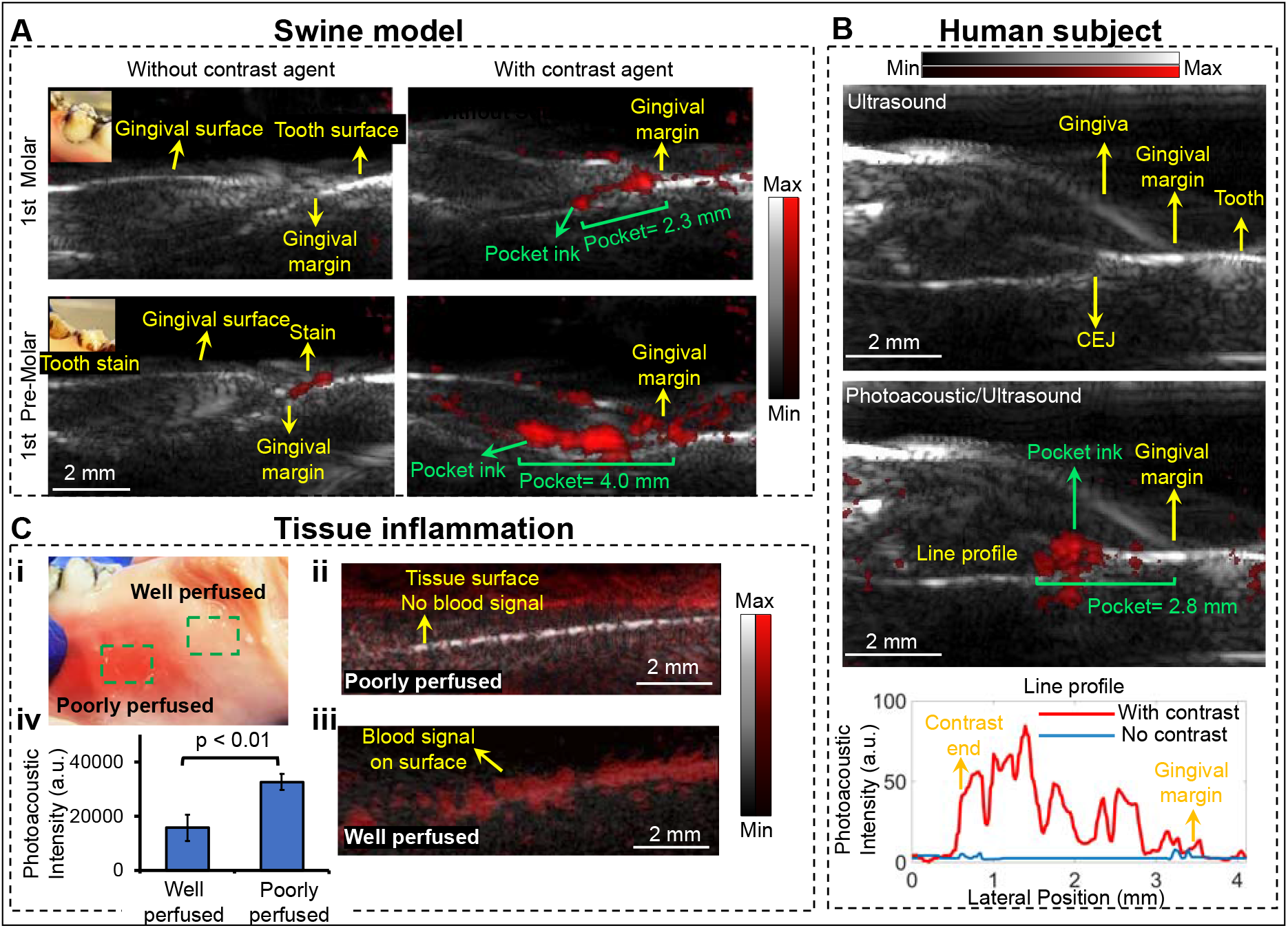
Periodontal pocket and gingiva inflammation imaging. A) Pocket depth measurement of two swine teeth before and after using the contrast agent. The insets show stains on tooth. Photoacoustic image in red scale is overlaid to ultrasound image in gray scale. B) Pocket depth measurement of 1^st^ molar #14 of a periodontal patient. The top panel is an ultrasound image of the molar. The middle panel is the photoacoustic/ultrasound image after applying the contrast agent. The bottom panel shows the line profile of the photoacoustic intensity in the pocket. C) Inflammation imaging. Panel i shows the swine gingiva with well-perfused tissue and poorly perfused tissue. Panel ii shows a photoacoustic/ultrasound image of the poorly perfused tissue. Panel iii shows photoacoustic/ultrasound image of the well-perfused tissue. Panel iv is the statistics that shows the overall photoacoustic intensity of healthy tissue (poorly perfused) and inflamed tissue (well perfused).

### 3.1. Device

The periodontal imaging system includes a photoacoustic/ultrasound imaging transducer, a laser-diode module, and a research-based ultrasound DAQ system (see Methods for configurations) (**Fig. 1(A)**). Photoacoustic imaging requires optical excitation, and the transducer integrates a transparent window and mirror for light delivery for photoacoustic imaging. This device uses a laser-diode module that delivers light to the transducer via an optical fiber. The transducer receives the photoacoustic and ultrasound signals which are processed by the DAQ system. A supplementary video is shared online that shows the operation of the system in periodontal pocket imaging (see **SM1** in Supplementary Information).

The laser-diode module contains a pulsed laser diode, a function generator board, coupling lens, and a heat sink (**Fig. 1(B)**). The laser-diode module’s dimensions are 35 cm (length) × 19 cm (width) × 28 cm (height), and it weighs ∼5 kg. The laser diode generates 1-KHz laser pulses which are coupled into the fiber and delivered remotely to the transducer for photoacoustic imaging. Thus, the transducer side (**Fig. 1(C)**) does not have any light source or thermal system making the transducer more compact than that of LED-based system. Alternative LED-based photoacoustic imaging systems require the LED arrays and heatsink to be assembled on the transducer which increase the size^13^. The laser-diode module also provides trigger in & out channels through a function generator board for synchronization. The laser-diode module is also more compact than a regular Q-switch laser, which makes it suitable for chairside imaging^14^. One limitation of the laser diode is that the pulse width is 100 ns, and thus it can only generate photoacoustic signal below 10 MHz^15^. Thus, we used the lower half of the transducer bandwidth for photoacoustic imaging and its full bandwidth for ultrasound imaging.

The transducer integrates a 19-MHz ultrasound transducer (average -6dB bandwidth of 48.9%) facing towards the tissue while the handle can project out from the gum parallel to the tissue surface (**Fig. 1(C)**). This toothbrush-shaped design can image the posterior teeth in contrast to conventional transducers^16, 17^. The transducer also integrates a transparent window and mirror for light delivery for photoacoustic imaging.

### 3.2. Characterization

We characterized the light homogeneity, light stability, penetration depth, and resolution of the system for photoacoustic imaging. The periodontal pocket and its associated tissue are less than 10 mm across the gingiva and tooth, and less than 4 mm deep from the gingiva surface. An array of pencil leads (10 mm wide, 5 leads) was put in a phantom solution (see Methods) 8 mm below the transducer to examine the light homogeneity. Pencil leads are strong light absorber, and their photoacoustic intensity correlates well with the local light intensity^18^; stronger photoacoustic signal indicates stronger light focus. All five pencil leads are distinguishable in the 10-mm range in the photoacoustic image with similar intensity (**Fig. 2(A)). Fig. 2(A)** shows the statistics that the photoacoustic intensity of the 5 pencil leads have only 11% relative standard deviation (RSD), which indicates uniform illumination.

We also measured the stability of the laser diode by monitoring its intensity in one hour duration (**Fig. 2(B)**). The light intensity dropped from 0.74 mJ/cm^2^ to 0.58 mJ/cm^2^ in the one hour with 20% variation. Three polyethylene tubes loaded with 20 μL contrast agent (cuttlefish ink solution) were fixed by a 3D printed holder. They were put in the phantom at different depths (7 mm, 11 mm, and 15 mm) from the transducer. The first two tubes are distinguishable while the third is barely observable with photoacoustic imaging (**Fig. 2(c)**). **Fig. 2(C)** shows that the signal-to-noise ratio (SNR) decreases with imaging depth which is expected due to light attenuation. These results demonstrate that the system can image contrast agents as deep as 11 mm in tissue with more than 10-dB SNR. The pulse energy of the laser diode is sufficient for periodontal pocket imaging (less than 4 mm deep).

The lateral and axial resolution of the system in photoacoustic imaging were determined by imaging the cross-section of a human hair (100-µm diameter). We defined the FWHM (full width at half maximum) of the lateral and axial amplitude distributions as the axial and lateral resolution, respectively^19^. The lateral and axial amplitude distributions across the hair were extracted (**Fig. 2(D)**), and the lateral and axial resolution in photoacoustic mode were 192 µm and 352 µm, respectively. The lateral and axial resolution values in ultrasound-only mode were 142 µm and 102 µm by imaging a 30-µm nichrome wire.

### 3.2 Ultrasound imaging of swine and human teeth

We next evaluated the performance of the transducer in tooth in ultrasound mode. One swine tooth and two human teeth were imaged *ex vivo* and *in vivo*, respectively. Swine teeth are a common model because they have a similar structure as human teeth^20^. The transducer resolves the structures in both the swine tooth imaging (1^st^ molar, **Fig. 3(a))** and the human tooth imaging (Cuspid #6, **Fig. 3(b)**) including gingival surface, tooth surface (occlusal surface), gingival margin, alveolar bone, and alveolar bone crest (ABC). The tooth surface has stronger ultrasound intensity because it has higher impedance than other tissue. Moreover, the cementoenamel junction (CEJ) is distinguishable in the human tooth (**Fig. 3(b))**. Resolving these structures and their positions are very helpful to the diagnosis of periodontal diseases^12^. Moreover, we imaged the 1^st^ molar of a human subject (**Fig. 3(c)**) *in vivo*. Those structures labeled in **Fig. 3(a)** and **3(b)** are also observable in **Fig. 3(c)**, which demonstrated that the toothbrush-shaped transducer can operate in the posterior area to image molars *in vivo*, which could be a big challenge for other ultrasound transducers.

We performed photoacoustic/ultrasound dual-mode imaging to resolve the periodontal pockets both *ex vivo* and *in vivo*. We used a cuttlefish ink solution as photoacoustic contrast agent which highlights the pocket area. The ultrasound image in grayscale shows the tooth structure, and photoacoustic image in red scale reveals the contrast agent i.e., the periodontal pocket. Photoacoustic/ultrasound images are overlaid together. Two swine teeth (1^st^ pre-molar and 1^st^ molar) were imaged *ex vivo* (**Fig. 4(A)**) and a molar of a periodontal patient was imaged *in vivo* (**Fig. 4(B)**). **Fig. 4(A)** shows the photoacoustic/ultrasound images of the two swine teeth before applying the contrast agent (left panels) and after (right panels). Clearly, there is no photoacoustic signal in the periodontal pocket before applying the contrast agent (Left panels). The photoacoustic signal from the 1^st^ pre-molar image was caused by the tooth stain as shown in the inset photo. After applying the contrast agent, the contrast agent is seen as a line below the gingival margin, i.e., the periodontal pocket. This confirms that the photoacoustic transducer can detect the periodontal pocket *ex vivo*. By drawing a line between the gingival margin in the ultrasound image and the contrast end in the pocket in the photoacoustic image, we measured the pocket depth to be 2.3 mm for the 1^st^ molar and 4.0 mm for the 1^st^ pre-molar, which is close to 2 mm and 4 mm respectively by using a clinical periodontal probe.

We further imaged the 1^st^ molar #14 of a periodontal patient (**Fig. 4(B)**). The diseased patient was diagnosed with stage 2, grade B periodontitis. Tooth #14 was characterized via conventional probing. The probing pocket depth (PPD) depths were (from distal to medial to mesial) 4-3-5 mm on the buccal side and 4-3-6 mm on the lingual. The Clinical attachment level (CAL) buccal (from distal to medial to mesial) was 4-3-5 mm, and the CAL lingual was 4-3-6 mm. Similar to the swine teeth, the photoacoustic signal shows up in the pocket after applying the contrast agent (**Fig. 4(B)**). **Fig. 4(B)** also shows the line distributions of the photoacoustic signal across the pocket. As can be seen, the pocket area has much stronger photoacoustic intensity (Red line) after applying the contrast agent. The pocket depth is measured as 2.8 mm (medial, buccal side). Importantly, our imaging assessment of the probing depth occurred medially is within 7% of the value (3 mm) determined by using a periodontal probe invasively. The results demonstrate that the transducer can image the periodontal pocket of molars *in vivo*.

Besides periodontal pocket, gingival inflammation also occurs with periodontal disease^21^. It is expected the inflamed tissue is well perfused and hence has more blood signal than healthy tissue (poorly perfused), which can be detected by imaging the hemoglobin by photoacoustic imaging. We performed photoacoustic imaging of the well perfused tissue and poorly perfused (healthy) tissue of the swine tooth model, separately (**Fig. 4(C). i**). As can be seen, well perfused tissue has much stronger PA signal close to the gingiva surface (**Fig. 4(C). ii)**) than poorly tissue (**Fig. 4(C). iii)**). The statistics in **Fig. 4(C). iv** shows that the overall photoacoustic intensity of the well perfused tissue is around two times higher than the poorly perfused tissue.

## 4. Conclusion

In conclusion, we evaluated a tooth-brushed-shaped photoacoustic/ultrasound transducer for full-mouth periodontal pocket imaging. The unique tooth-brush design allows the transducer to fit in the posterior area of gum and image molars. The device uses a laser diode as the light source to reduces the costs. We performed *ex vivo* swine teeth imaging and *in vivo* periodontal patient imaging. The results demonstrated the transducer can image the periodontal pocket in molars *in vivo*.

## 5. Back matter

### 5.1 Funding

A.H. and R.K. acknowledge R43 DE031196. J.V.J acknowledges R21 DE029025 and UL1 TR001442.

### 5.2. Disclosures

R.K., A.H., and J.V.J. are co-founders of StyloSonic, LLC.

### 5.3 Data availability

Data can be requested from the authors at any time. A supplementary video is shared in the supporting information that shows the operation of the system in periodontal pocket imaging.

